# The impact of inhibitor size and flexibility on the binding pathways to c-Src kinase

**DOI:** 10.1101/2022.10.25.513784

**Authors:** Ai Shinobu, Suyong Re, Yuji Sugita

## Abstract

Considering dynamical aspects of protein-drug binding processes is inevitable in current drug compound design. Conformational plasticity of protein kinases poses a challenge for the design of their inhibitors, and therefore, atomistic molecular dynamics (MD) simulations have often been utilized. While protein conformational changes have been increasingly discussed, a fundamental yet non-trivial question remains for the effect of drug compound flexibility, which is hardly detectable from experiments. In this study, we apply two-dimensional replica-exchange MD simulations as enhanced sampling to investigate how c-Src kinase can bind PP1, a small inhibitor, and dasatinib, a larger inhibitor with greater flexibility. 600 microseconds simulations in total sample binding and unbinding events of these inhibitors much more frequently than conventional MD simulation, resulting in statistically converged binding pathways. While the two inhibitors adopt a similar mechanism of multiple binding pathways, the non-canonical binding poses become less feasible for dasatinib. A notable difference is apparent in their energetics where dasatinib stabilizes at intermediate states more than PP1 to raise the barrier toward the canonical pose. Conformational analysis shows that dasatinib adopts linear and bent forms for which relative populations are altered upon binding. We further find hidden conformations of dasatinib at intermediate regions, and unexpectedly one of them could efficiently bypasses the intermediate-to-bound state transition. The results demonstrate that inhibitor size and flexibility impact the binding mechanism, which could potentially modulate inhibitor residence time.

## Introduction

The development of small organic kinase inhibitors has greatly contributed to the advance in cancer therapy for the past decades^1, 2^. Currently, over 70 approved drugs are available for cancer treatment, while their effectiveness is severely hampered by drug resistance as well as by off-target effects^3^. Protein kinases are the enzymes that catalyze the transfer of a phosphate from ATP to a target protein. Various cellular mechanisms, including signal transduction rely on the proper functions of kinases, thus their dysregulation can result in cancer or other diseases. Kinases are considered as difficult targets for specific inhibitor design due to their conformational plasticity, high similarity of the active site (ATP binding site), and frequent occurrence of mutations. Thus, rational design methods that go beyond the traditional approach, which are based on stable kinase-inhibitor complexes are attractive.

In the past decades, dynamical aspects of kinase-inhibitor binding have been increasingly utilized for inhibitor design. Exploiting various conformational states of kinases^4, 5^, which are regulated by key structural elements, such as the Gly-rich or phosphate binding loop (G-loop or P-loop), an αC-helix, and an activation loop (A-loop) (Figure 1A), have contributed to the development of conformation-selective and allosteric inhibitors targeting distinct conformations and/or sites other than the commonly targeted active states^6, 7^. Another approach for inhibitor design is the exploration of inhibitor (un)binding pathways, which allow to utilize not only the binding affinity but also kinetic factors such as residence time. Recent *in-cell* screening of the binding of imatinib against Abl kinase mutants showed that many mutations affect the dissociation kinetics rather than the binding affinities^8^. NMR and other fast kinetics experiments proposed a binding model consisting of fast binding followed by a slow induced-fit process as the determinant^9^, although the atomistic details of the dynamic nature of kinase-inhibitor interactions are hardly accessible from experiments alone.

**Figure 1.**
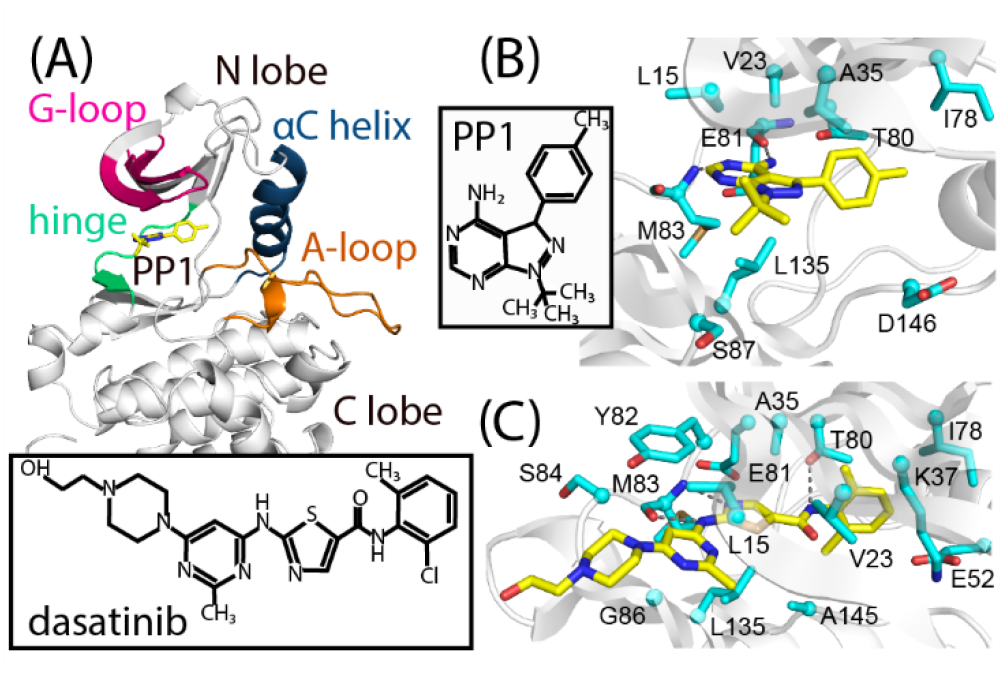
Structures of the c-Src-PP1 and c-Src-dasatinib complexes. (A) The structure of c-Src-PP1 from modeled X-ray structures (PDB IDs: 1Y57^25^/1QCF^26^). (B), (C) Binding sites of c-Src-PP1 (B) and c-Src-dasatinib (C) from X-ray structures (PDB ID: 1Y57^25^/1QCF^26^ and 1Y57^25^/3G5D^27^, respectively). PP1 and dasatinib are shown in yellow, interacting residues are shown in cyan. Chemical structures of PP1 and dasatinib are shown in (B), (C).

Molecular Dynamics (MD) simulations with enhanced sampling techniques have greatly advanced our understanding of inhibitor (un)binding pathways^10-12^. Enhanced sampling MD methods can realize a considerably larger number of (un)binding events than conventional MD simulations, allowing statistically reliable characterizations of (un)binding processes. Such simulations have been used to explore kinase conformational dynamics^13-15^ as well as for establishing relations between inhibitor-binding and observed kinetics^16-24^. These computational studies targeted various inhibitors including the well-known cancer drug imatinib, and suggested that the inhibitor can (un)bind through multiple routes by transiently interacting with key structural elements regulating kinase conformations such as the G-loop or the αC-helix.

We previously^28^ analyzed the binding of PP1 (an ATP competitive inhibitor) to c-Src kinase (Figures 1B and S1) using the two-dimensional (2D) replica-exchange method— gREST/REUS, a combined replica-exchange umbrella sampling (REUS^29, 30^) and generalized replica exchange with solute tempering (gREST^31^). In the study, we characterized multiple binding poses and pathways based on the free-energy landscape, which was obtained from extensive MD trajectories with 100 (un)binding events during the simulation. The results showed that modulation of transient intermediates, such as the encounter complex, is potentially effective for optimizing the drug-target residence time. However, PP1 (MW of 281) is a small compound, compared to many approved inhibitors which have molecular weights (MWs) of more than 500 and are accompanied by additional internal degrees of freedom. This raises the question of whether the increase in molecular size and flexibility changes the fundamental reasoning for rational compound design. Here, we investigate the binding of dasatinib (a cancer drug sold as Sprycel, Bristol-Myers Squibb, MW of 488)^32^ to c-Src kinase (Figures 1C and S1) to answer this question. We find that PP1 and dasatinib both bind to their binding sites through multiple routes but have different energetics due to the increased flexibility of dasatinib.

## Results

### Two inhibitors adopt similar binding pathways

We examine the overall binding processes of the two inhibitors using gREST/REUS simulations. Figures 2A (PP1) and 2B (dasatinib) show the locations of the inhibitor with respect to the protein at 310 K. The simulations were initiated from both the bound and unbound states (referred to as “forward” and “reverse”, respectively). Both PP1 and dasatinib bind non-specifically to various sites at the N-lobe of c-Src including the back side of the binding pocket (cyan colored regions), while the distribution of dasatinib is slightly broader than for PP1. In the current work, we focused on exploration of the N-lobe region, but we expect similar nonspecific bindings in the C-lobe region^33, 34^. As the inhibitor approaches the binding pocket (yellow-colored regions), the distribution is restricted to the interface region between the N- and C-lobes. The inhibitor is located around the pocket with populations near both the hinge and the G-loop, suggesting two alternative binding pathways.

**Figure 2.**
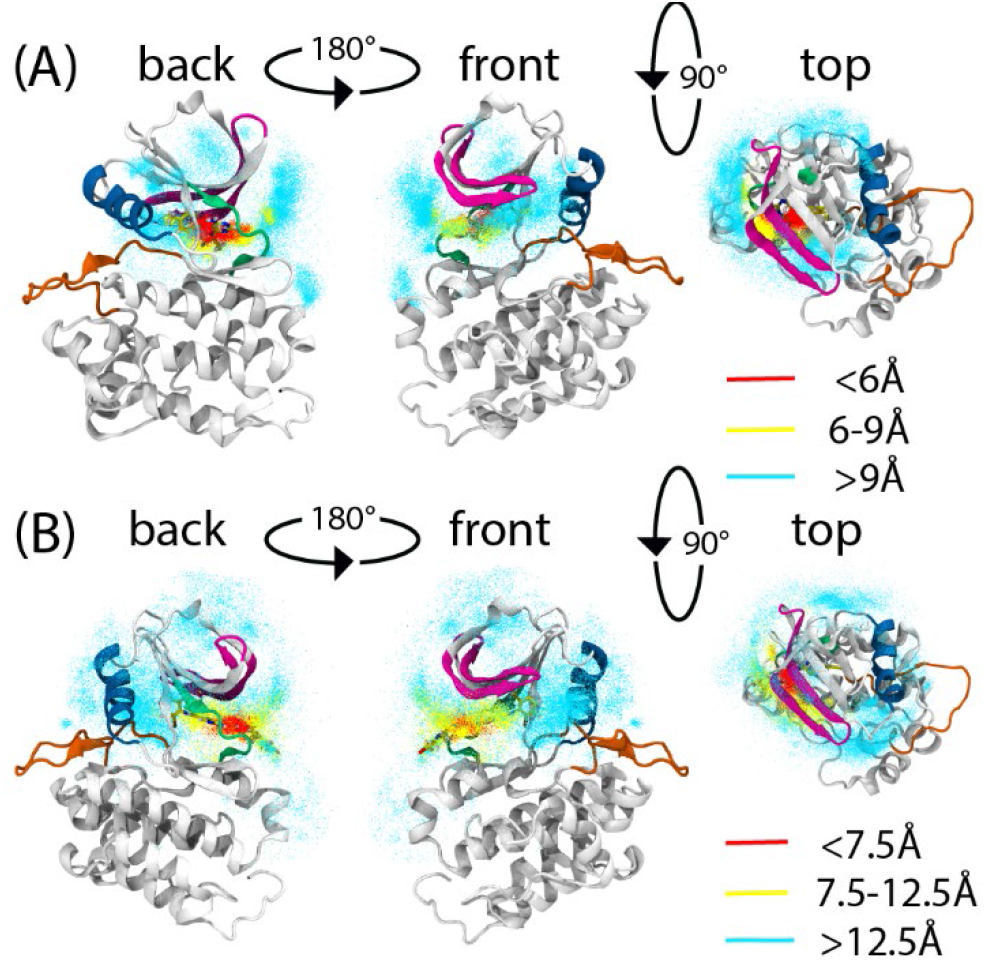
Location of the inhibitor with respect to c-Src kinase for PP1 (A) and dasatinib (B) in the gREST/REUS simulations at 310 K (for both “forward” and “reverse” simulations). Inhibitor locations indicate the coordinates of the C9 atom for PP1 and the C11 atom for dasatinib. Points are colored according to the protein-inhibitor distance.

Free-energy profiles along the kinase-inhibitor distance reflect the difference in the observed binding affinities of PP1 (IC^50^ of 170 nM^35^) and dasatinib (*K*_d_ of 11 nM^27^) as shown in Figures 3A (PP1) and 3B (dasatinib) (data from the “forward” simulations. See Figure S2 for the “reverse” simulations). For both systems, the position of the minimum free-energy agrees well with the kinase-inhibitor distance in the X-ray structures (PDB 1QCF^26^ for PP1 and PDB 3G5D^27^ for dasatinib). The difference in the binding free-energies between PP1 and dasatinib is roughly 2 kcal/mol, which reasonably agrees with the estimated value from the observed affinity (1.7 kcal/mol)^27, 35^. Since our simulations cover only half of the binding process (starting from the encounter state), the contribution from the initial association (approximately 6.5 kcal/mol^9^) should be added for estimating the absolute binding free-energy. For the subsequent analyses, we defined three regions of bound (“B”), intermediate (“I”), and encounter (“E”) according to the kinase-inhibitor distance: < 6.0 Å (B), 6.0-9.0 Å (I), >9.0 Å (E) for PP1 and < 7.5 Å (B), 7.5-12.5 Å (I), >12.5 Å (E) for dasatinib.

**Figure 3.**
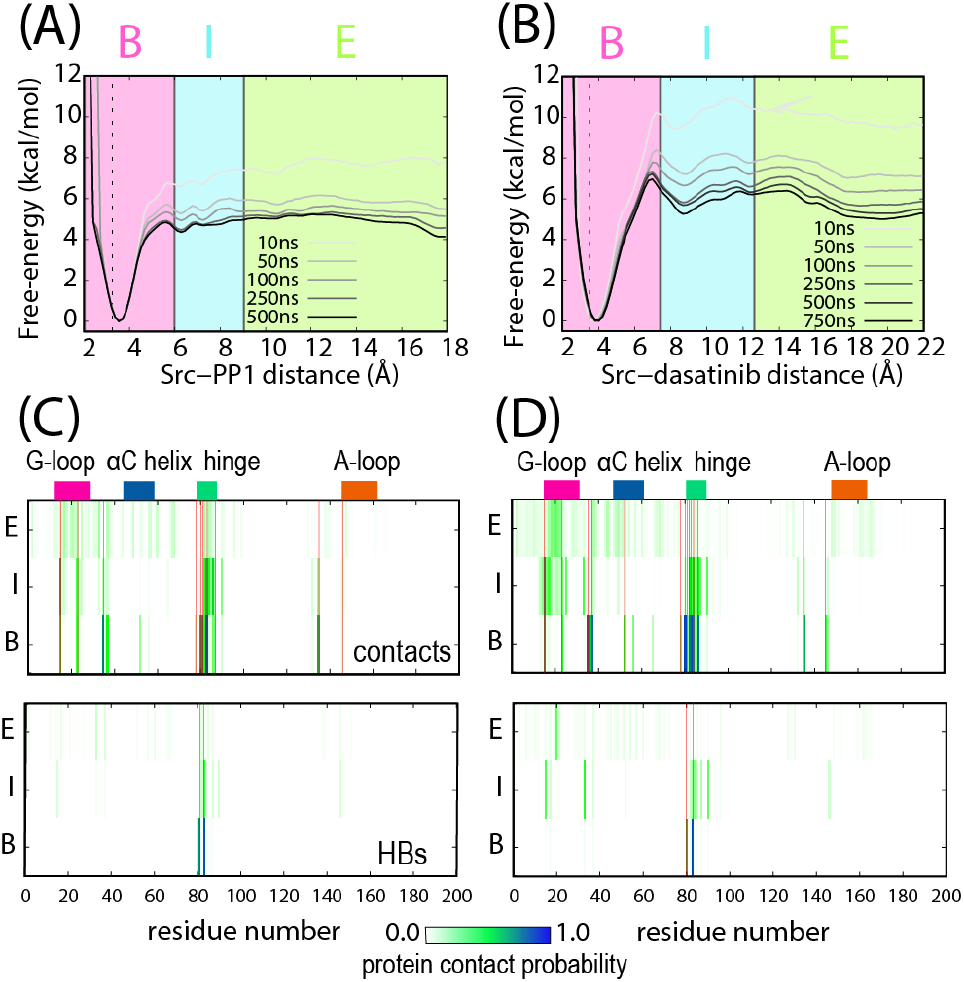
(A), (B) Free-energy profiles at 310 K along the c-Src-PP1 (A) and c-Src-dasatinib (B) distance. The kinase-inhibitor distances in the X-ray structure (3.25 Å, PDB 1Y57^25^/1QCF^26^ for PP1 and 3.48 Å, PDB 1Y57^25^/3G5D^27^ for dasatinib) are shown in dotted lines. Divisions between regions are marked in solid lines. (C), (D) Protein-inhibitor nonbonded contact (top) and HB (bottom) probabilities for c-Src residues 1-200 at 310K for PP1 (C) and dasatinib (D). Locations of the residues with which the inhibitor interacts in the X-ray structure are shown in red vertical lines.

### Dasatinib binding is accompanied by a free-energy barrier

Noteworthy, upon moving from the intermediate (I) to the bound region (B), there is a free-energy barrier for dasatinib which does not exist in the case of PP1. One of the factors which might contribute to the barrier is the increase in the number of interactions that dasatinib forms with c-Src surface residues due to its molecular size. Figures 3C (PP1) and 3D (dasatinib) show nonbonded contact and hydrogen bond (HB) probabilities between the inhibitor and kinase residues at the three regions, E, I, and B (see Figure S3 for the interaction sites on the c-Src three-dimensional (3D) structures, Tables S2 (nonbonded contacts) and S3 (HBs) for the interaction probabilities). For both PP1 and dasatinib, the kinase-inhibitor interactions are non-specific in region E, while they gain specificity in regions I and B. In region E, the inhibitor interacts with various sites on the N-lobe including those remote from the binding site such as the A-loop. Note that the interaction probability with the G-loop is slightly higher for dasatinib. On the other hand, the interactions with the G-loop and the hinge residues dominate in region I. The interacting residues include Leu15 and Gln17 in G-loop and Tyr82, Met83 (HB), Ser84 (HB), and Gly86 in hinge. Finally, the canonical bound pose is stabilized by strong HBs with the hinge residues (Glu81 and Met83 for PP1, and Thr80 and Met83 for dasatinib) as well as by other nonbonded interactions with Leu15, Ala35, and Leu135. Given that dasatinib has a larger molecular size than PP1, the number of nonbonded contacts and HBs for dasatinib in region I is greater, as shown in Figure 3. These interactions must be broken upon dasatinib entering the binding pocket, while raising the free-energy barrier.

### Transitions between states become difficult for dasatinib

Next, we characterize the binding pathways based on the free-energy landscape (FEL). A 2D FEL is constructed along the kinase-inhibitor distance as well as the inhibitor orientation, as we did in our previous study on PP1^28^ (Figures 4A and 4B). In brief, the inhibitor orientation (β) was defined as the angle formed between two bonds, L1-L2 from the inhibitor and P1-P2 from c-Src (Figure 4B, see supporting text for the full definition). Representative c-Src-inhibitor complex structures were obtained from *k*-means clustering analysis and their positions were marked on the FELs (see supporting text for detail on clustering).

**Figure 4.**
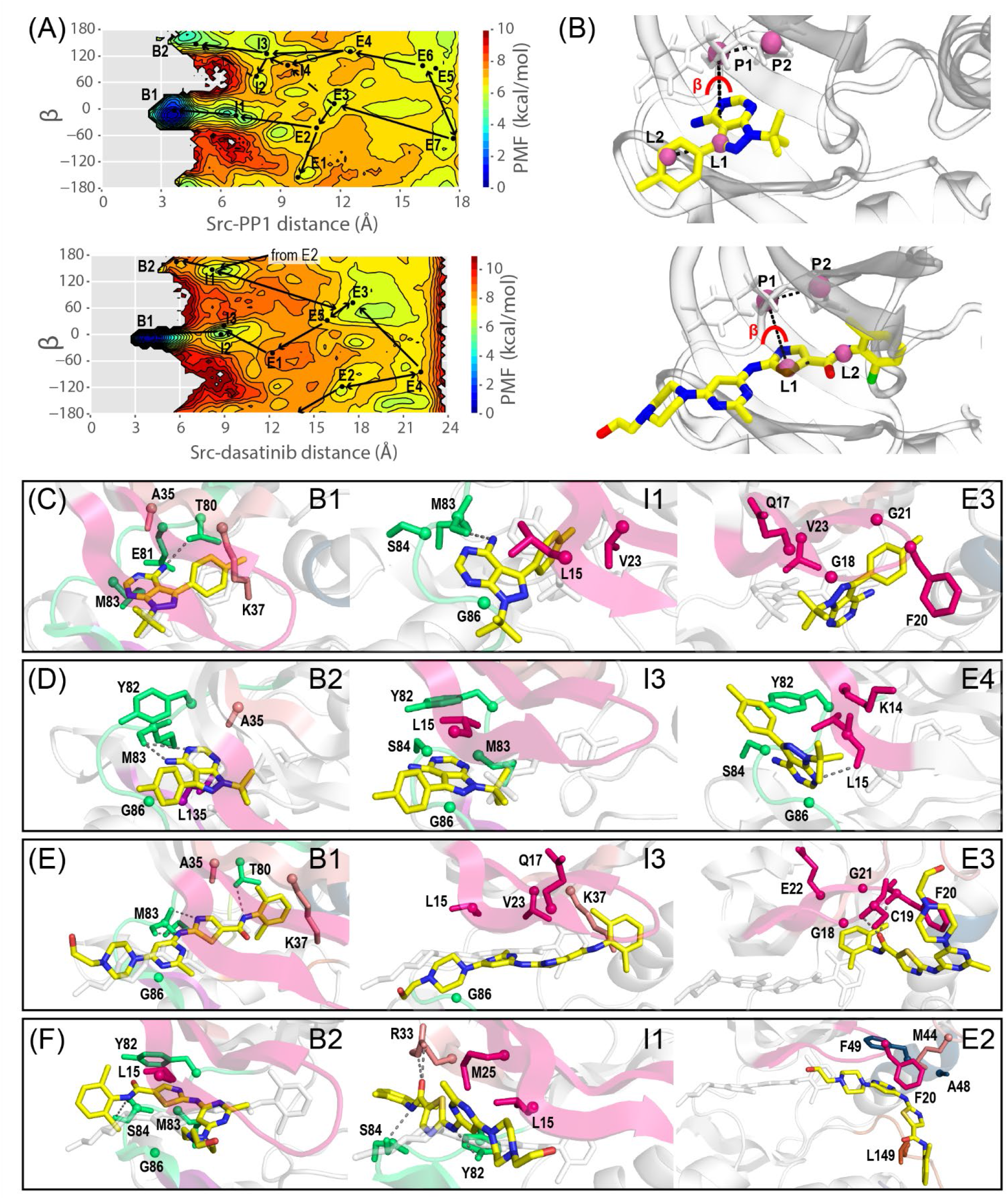
(A) The 2D free-energy landscapes along the kinase-inhibitor distance and the angle β at 310 K for PP1 (top) and dasatinib (bottom). (B) The definition of the angle β. Anchor points are shown in pink balls. Representative structures at 310 K for PP1 (C, D) and dasatinib (E, F). The inhibitors are shown in yellow. X-ray positions and structures of the inhibitors are shown in white. Five most interacting protein residues with the inhibitors are shown in licorice representation and colored according to the structural motif to which they belong. The Cα atoms of the residues are shown as spheres. Additional interacting residues in cartoon representation are colored. HBs are shown in dashed lines.

For both PP1 and dasatinib, the FELs are relatively flat in region E, while multiple states become separated by high energy barriers as the binding proceeds toward the I and B regions. The two bound poses, which differ in the inhibitor orientation (major canonical pose B1 and minor semi bound pose B2), are commonly identified on the FEL for both PP1 and dasatinib. The two bound states are well separated by a high free-energy barrier, and thus direct passage between the states is not likely. Namely, the distinct interactions in region E determine the final pose in region B ^28^. Possible routes connecting the E and B regions were drawn on the FELs, indicating that the E3 state is crucial for the canonical bindings for the two inhibitors. Notable difference is that the minor semi bound pose becomes less feasible for dasatinib. More importantly, dasatinib binding is accompanied by higher energy barriers than PP1 binding. Indeed, each of the three states, E3, I3, and B1, along the dasatinib major binding pathway is more distinctly separated by free-energy barriers than those along the corresponding pathway for PP1 (E3, I1, and B1).

The representative changes in the kinase-inhibitor interactions along the major and minor pathways are illustrated in Figures 4C-4F. We find a common feature that, in the major pathway toward B1, the inhibitor initially interacts with G-loop residues in the E3 states for both PP1 and dasatinib and then switches to interact with the hinge residues on its way to the canonical bound pose (Figure 4C for PP1 and Figure 4E for dasatinib). On the other hand, in the minor pathway toward B2, the interactions with the hinge residues dominate throughout the whole binding pathway (Figure 4D for PP1 and Figure 4F for dasatinib). Despite the above commonality, the increased molecular size of dasatinib increases the number of interactions, making transitions between states more difficult than in PP1. In addition, the inversion in orientation leads to a drastic change in the number and type of interactions, likely making the minor pose much less feasible for dasatinib than for PP1. Noteworthy, dasatinib changes its molecular shape along the binding pathways, which is less pronounced for PP1.

Here we demonstrated the parallel binding pathways on the selected coordinates, the kinase-inhibitor distance as well as the inhibitor orientation (β) for the “forward” simulations. The pathways that were identified on the other positional and orientational angles and the FELs for the “reverse” simulations are also provided as Figures S4 and S5, respectively, in the supporting information.

### Inhibitor flexibility could play a passive role in binding

PP1 has only one rotatable bond between the pyrazole and phenyl rings and adopts mostly a slightly tilted shape as seen in the representative poses (Figures 4C and D). In contrast, dasatinib has six rotatable bonds (φ_1_ - φ_6_, Figure S6) in the main frame^36^. Here we focus on φ_2_ - φ_4_ which affect the overall conformation of dasatinib. Dasatinib can adopt several bent forms in the I and E regions (Figures 4E and F), while in the canonical binding pose it adopts a linear conformation. To analyze the impact of conformational flexibility of dasatinib on the binding mechanism, we extract the representative conformers at each of the regions, E, I, and B, as well as those in solution (Figures 5A and 5B). For that purpose, we independently performed temperature replica-exchange MD (T-REMD)^30^ simulations of dasatinib in solution (see supporting text for details of the simulation). *k*-means clustering analysis shows that dasatinib adopts four main conformers (Figure 5B), which vary in the dihedral angles φ_2_ and φ_4_, while φ_3_ always maintains the same value. The linear conformer (I) is the X-ray bound conformer while the other conformers are bent and differ from their X-ray structure values in φ_2_ (II), φ_4_ (III), or both (IV).

**Figure 5.**
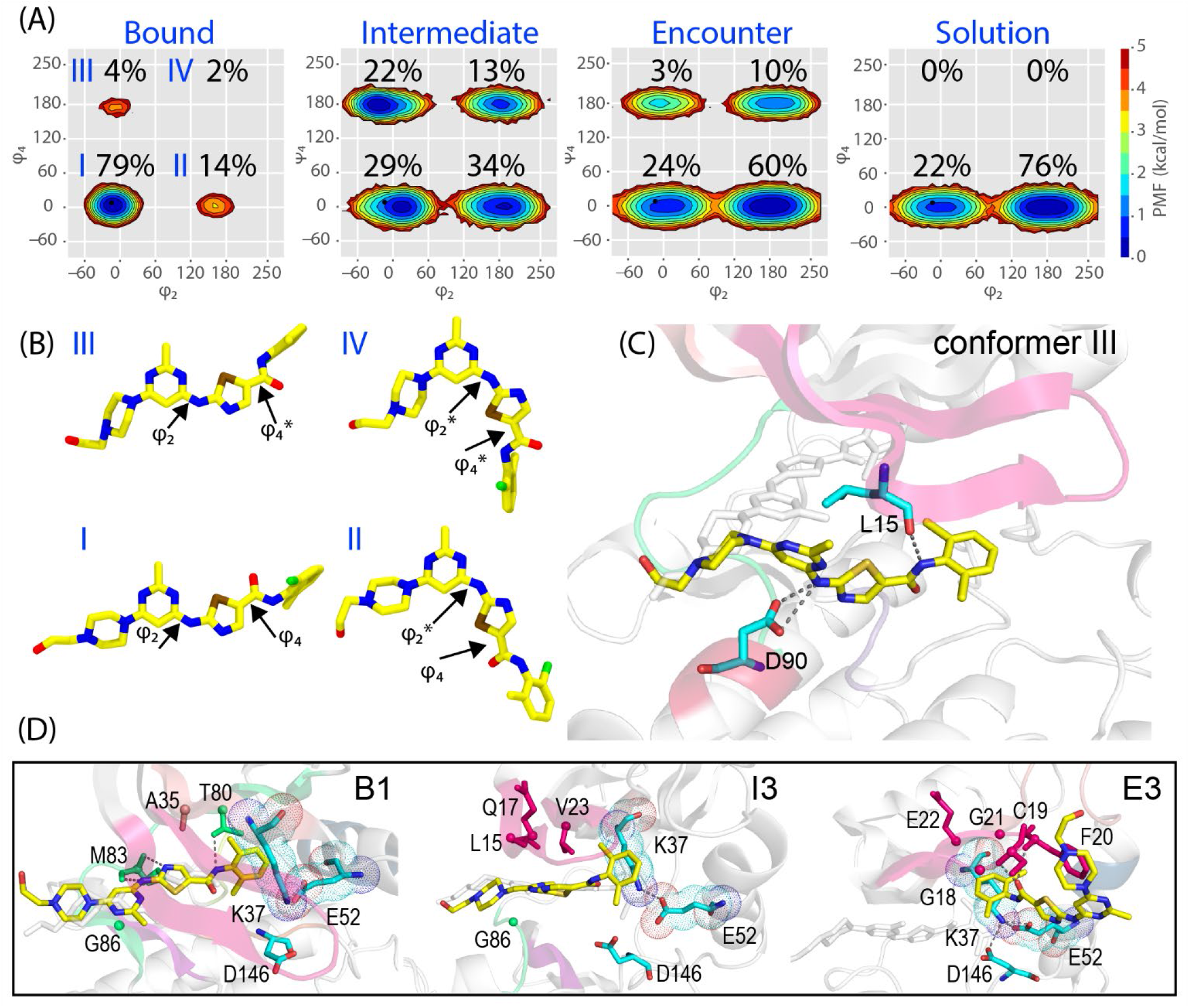
(A) The 2D Free-energy landscapes along the angles φ_2_ and φ_4_ for the three kinase-inhibitor distance regions and for dasatinib in solution at 310 K. Population at each conformer is written as percentage on the plots. (B) Most common conformers extracted from *k*-means clustering of dasatinib (number of clusters = 10). The angle differing from the X-ray structure value is marked in asterisk for each conformer. (C) Representative structure of dasatinib conformer III in the intermediate region (region I) extracted from *k*-means clustering for dasatinib at 310 K. Dasatinib is shown in yellow and the protein residues forming HBs with it are shown in cyan. Other interacting regions are colored. X-ray inhibitor position is shown in white licorice representation. (D) Representative structures at 310 K for dasatinib along the main pathway leading to B1. Dasatinib is shown in yellow, the X-ray location of dasatinib is shown in white. The five most interacting protein residues with the inhibitor are shown in licorice representation and colored according to the structural motif to which they belong. Additional interacting residues in cartoon representation are colored. HBs are shown in dashed lines. Residues Lys37 and Glu52 forming a salt bridge are shown in cyan.

The relative population of the four conformers I-IV changes along the binding process (Figure 5A). When dasatinib is free in solution, only the linear (I) and one bent conformer (II) exist with the bent form dominating (76%). In the E region, where dasatinib resides on the protein surface while forming mostly non-specific interactions with the surface residues, the other two bent forms (III and IV) appear. In the I region, where the inhibitor interacts mainly with the G-loop and the hinge residues, the population of the latter two bent forms increase, mostly for conformer III. Upon entering the binding pocket, the population of conformers III and IV almost vanish, and the linear form (I) becomes dominant with 79% of the population. The hidden conformers III and IV that appeared only in the I and E regions but not in solution or in the B region can be understood by examining the representative poses in surface representation (Figure S7A). While in the E and I regions, dasatinib forms transient interactions with surface residues of c-Src by flexibly adapting its shape to the protein surface. The bent conformer will seemingly increase the residence time of the inhibitor on the protein surface. On the other hand, in the B region, it must exclusively adopt the linear conformation (I) to maximize the interaction energy, which is accompanied by an entropic penalty. This penalty is likely one of the contributors to the free-energy barrier that is observed only for dasatinib.

To confirm whether there is an active mechanistic role to the bent conformers in the E and I regions, we look at their individual contact maps (Figure S7B). In the E region, conformers I and II tend to form interactions with the G-loop and the A-loop while conformers III and IV tend to form interactions with the αC-helix. In the I region, it is notable that conformer III forms strong and sharp interactions with the G-loop and the αC-helix, two of these interactions are HBs with Leu15 and Asp90 (Figure 5C). In the major pathway for dasatinib binding (E3-I3-B1), a conserved salt bridge between Lys37 and Glu52 is seen intact. Dasatinib in E3 and I3 is outside the binding pocket (in front of the salt bridge), whereas, in B1, it is inside the pocket (behind the salt bridge). We suggest the following mechanism in which dasatinib adopts the bent conformer III and by forming the strong HBs with Leu15 and Asp90, manages to place itself at the binding pocket entrance without breaking the Lys37-Glu52 salt bridge. The free-energy barrier observed for dasatinib is thus composed of breaking the Leu15-Asp90 HB as well as the transition from the bent conformer III to the linear conformer I.

## Discussion

In this study, we investigated the binding of two small kinase inhibitors of different molecular size, PP1 and dasatinib, to c-Src kinase. Simulations of 600 microseconds (240 µs and 360 µs for PP1 and dasatinib, respectively) show an overall similarity between the inhibitors, with two parallel binding pathways among which the G-loop-bound encounter state serves as a main gate to the native pose. PP1 enters the pocket through transient cleavage of the conserved Lys37-Glu52 salt bridge and concomitant αC-helix outward rotation. Noteworthy, dasatinib is stabilized at intermediate states more than PP1 while raising the free-energy barrier toward the bound state. A recent simulation of imatinib binding to Abl kinase also showed the presence of various intermediates^37^ which could extend the residence time on the protein surface. These findings could account for, at least partly, the slow induced fit process which was proposed by a stop-flow experiment^9^. Given that dasatinib and imatinib have similar molecular weights, this view is likely applicable to many approved drug compounds with molecular weights above 500.

The conformational flexibility of protein kinases has been extensively studied^4, 5, 13-15^, while that of the inhibitors during the binding processes was rarely discussed in the past. Our simulations show that dasatinib adopts linear and bent forms whose relative populations are altered upon binding. A similar conformational switch has also been suggested for imatinib binding to Abl kinase^37^. Together with the significant reduction of the conformational flexibility upon binding^22^, such changes in inhibitor conformation can contribute to the free-energy barrier, both enthalpically and entropically. Intriguingly, we found hidden conformations of dasatinib in the intermediate region. One of these conformations could bypasses the main intermediate leading to the bound state transition, allowing dasatinib to enter the pocket while the conserved Lys37-Glu52 salt bridge remains nearly intact. This finding suggests an alternative view to that obtained from a previous simulation of dasatinib dissociation from c-Src kinase, which suggested that the salt-bridge must be cleaved in order for dasatinib to dissociate^16^. Our simulations, which extensively sample the inhibitor and kinase binding site residues, provide a statistically converged description for both binding and unbinding pathways. Closely concordant pathways have also been reported for dasatinib binding using unbiased simulations^33,38^. Traditionally, properties such as conformational flexibility and the number of rotatable bonds have been considered in relation to oral bioavailability^36^. The flexibility of inhibitor conformation may also be considered for optimizing the binding process and associated kinetic properties such as residence time.

Drug resistant mutations pose an inevitable difficulty in the development of kinase inhibitors^39, 40^. Generally, kinases develop resistance through either orthosteric or allosteric mutations. Recent profiling study on dasatinib resistance in c-Src kinase showed that mutations in either the inhibitor contact residues (Leu15, Val65, Tyr82, and Thr80) or those proximate to the G-loop region (Glu12, Val13, Lys14, Glu22, and Trp24) could develop the resistance^41^. For example, a mutation at the well-known gate keeper residue, T80M or T80I, abolishes dasatinib activity by sterically hindering the binding pocket^42^. Our simulations suggest that mutations at Leu15, Glu22, and Try24, located next to the G-loop, could affect the binding process, likely contributing to the resistance. The latter group mutations have been considered to hinder the autoinhibitory SH2/SH3 interaction and increase the phosphotransferase activity by modulating the conformational dynamics of the N-terminal lobe including the G-loop^41^. We speculate that those mutations could also modulate the (un)binding pathways of dasatinib to affect the residence time, as reported for imatinib dissociation from Abl kinase^24^. Further exploration will lead to understanding the resistance mechanism and could assist in designing inhibitors which can overcome the resistant mutations.

## Methods

### System preparation

The simulated systems were constructed in our previous works^28, 43^. Here we briefly describe the modeling procedure. For c-Src-PP1 we used only the kinase domain (residues 20-533, renumbered 2-275 in this work) from the X-ray crystal structure of active-like c-Src kinase (PDB ID: 1Y57^25^) and replaced the co-crystalized ligand with PP1 from PP1-bound Hck (PDB ID: 1QCF^26^). For c-Src-dasatinib, we used the kinase domain of c-Src kinase (PDB ID: 1Y57^25^) and replaced the co-crystalized ligand with dasatinib from dasatinib-bound c-Src (PDB ID: 3G5D^27^). We solvated the kinase-inhibitor complexes with water molecules (7,698 and 13,992 for c-Src-PP1 and c-Src-dasatinib, respectively) and added sodium counter ions (six ions for each of the systems) for neutralization. Modeling was performed with Amber-Tools16^44^. The systems were subsequently minimized for 1,000 steps while restraining protein backbone atoms by 10.0 kcal/mol/Å^2^. Then they were heated to 310 K under NVT conditions for 105 ps followed by equilibration in NPT for 105 ps. Finally, all restraints were removed, and the systems were equilibrated for 1.05 ns under NPT conditions.

### Two-dimensional gREST/REUS setup and simulation

We defined the solute region in gREST^31^ as the dihedral angle and the nonbonded energy terms of the ligand (“inhibitor”) and 12 binding site residues (residues defined as SITE residues according to the crystal structure 1QCF^26^). We determined the solute temperatures using the automatic parameter tuning tool in the GENESIS MD program^45, 46^. Eight gREST replicas were defined in the range of 310-692 K for c-Src-PP1 and 310-663 K for c-Src-dasatinib. The reaction coordinate for the REUS^29, 47^ dimension was chosen as the center of mass distance (“kinase-inhibitor distance”) between the inhibitor heavy atoms and the backbone atoms of residues Ala35 and Leu135 of c-Src. 30 REUS replicas of increasing kinase-inhibitor distance were prepared by stepwise pulling of the inhibitor from its bound position (“forward” direction) and subsequently pulling it back to its bound-conformation kinase-inhibitor distance (“reverse” direction) while restraining the protein Cα atoms to avoid its artificial deformation. REUS replicas cover the range of 3.0 Å to ∼18.0 Å and ∼23.0 Å for c-Src-PP1 and c-Src-dasatinib, respectively. The position of REUS replicas as well as the force constants of the umbrella potential were finetuned to achieve a satisfactory coverage of the REUS space. Two initial REUS kinase-inhibitor distance setups were prepared, from the “forward” and the “reverse” pulling simulations, a total of four simulation systems. Analysis presented in the main text refer to the “forward” simulations, unless specified otherwise. The full parameters used for the simulations are presented in Table S1. The detailed procedure for gREST solute temperature and REUS replica tuning was described in our previous paper^43^.

The gREST/REUS simulations were performed using the GENESIS MD program^45, 46^ (version 2.0 beta^48^). The AMBER ff99SB-ILDN^49, 50^ force field was used for the protein, GAFF^51^ (with AM1-BCC) was used for the inhibitors, and the TIP3P^52^ model was used for water molecules. The SHAKE^53^ algorithm was used for constraining bonds involving hydrogen atoms and the SETTLE^54^ algorithm was used for keeping water molecules rigid. The Particle-mesh Ewald^55, 56^ summation was used for evaluating long-range electrostatic interactions while the cutoff distance for non-bonded interactions was set to 8 Å. Simulations were performed in the NVT ensemble at 310 K using the Bussi thermostat^57^. The RESPA integrator^58^ with hydrogen mass repartitioning^59^ on solute atoms was used with a timestep of 3.5 fs.

A total of 240 (8×30) replicas were prepared for each system for the 2D gREST/REUS simulations. All replicas were first equilibrated for 1.05 ns, followed by production runs of 500 ns and 750 ns per replica (for c-Src-PP1 and c-Src-dasatinib systems, respectively). Exchanges were attempted each 2.1 ps alternatively in the gREST and the REUS dimensions. Frames for analysis were written every 10.5 ps. Simulations were performed on the supercomputer Fugaku^60^. GENESIS 2.0 beta^48^ was optimized for Fugaku for obtaining a speed of over 50 ns/day using 480 nodes.

## ASSOCIATED CONTENT

### Supporting Information

Supporting text: Definition of kinase and inhibitor relative position and orientation; Obtaining structural ensembles from clustering; T-REMD simulations of dasatinib in water; Supporting tables: S1: gREST/REUS simulation parameters; S2: Kinase-inhibitor interaction probabilities; S3: Kinase-inhibitor HB probabilities; Supporting figures: S1: Canonical binding pose interactions; S2: Free-energy profiles and contact maps for reverse simulations; S3: Mapping of Interacting protein residues; S4: Free-energy landscapes for forward simulations; S5: Free-energy landscapes for reverse simulations; S6: Definition of rotatable dihedral angles for dasatinib; S7: Representative inhibitor poses in surface representation, contacts maps for individual inhibitor conformers; (PDF); Supporting files: structures files of the most common poses along the binding pathways (PDB).

This material is available free of charge via the Internet at http://pubs.acs.org.

## AUTHOR INFORMATION

### Author Contributions

The manuscript was written through contributions of all authors. All authors have given approval to the final version of the manuscript.

### Notes

The authors declare no competing financial interest.

## ACKNOWLEDGMENT

This work was supported in part by MEXT JSPS Kakenhi (grant number 19H05645, 21H05249 (to Y.S.), 19K12229 (to S.R.)), RIKEN pioneering projects “Biology of Intracellular Environments”, and “Glycolipidologue Initiative” (to Y.S.), and MEXT program for promoting research on the supercomputer Fugaku (JPMXP1020200101, JPMXP1020200201 (to Y.S.)). The computer resources are provided by the HPCI system research project (Project ID: hp200129, hp200135, hp210172, hp210177, hp220164, and hp220170) and by RIKEN Advanced Center for Computing and Communication (for HOKUSAI BigWaterfall).

