## Supplementary information for "The impact of inhibitor size and flexibility on the binding pathways to c-Src kinase"

**Definition of kinase and inhibitor relative position and orientation**

We follow the definition of Wang et al.^1^ and Jo et al.^2^ (which was also used in our previous work.^3^) for describing the relative position and orientation of the inhibitor with respect to the protein kinase. Six anchor points (L1, L2, L3, P1, P2, P3) were defined where each anchor point was defined by the center of mass (COM) of a group of atoms as follows. L1 is the COM of the inhibitor atom L0 closest to the inhibitor COM and atoms bonded to L0. P1 is the COM of the kinase backbone heavy atoms of the residue closest to the COM of the kinase. P2 is the COM of the kinase backbone heavy atoms of the residue satisfying 30⁰ ≤ ∠L1P1P2 ≤150⁰. P3 is the COM of the kinase backbone heavy atoms of a residue satisfying 30⁰ ≤ ∠P1P2P3 ≤150⁰. L2 is the COM of an inhibitor heavy atom satisfying 30⁰ ≤ ∠P1L1L2 ≤150⁰ and the atoms bonded to the heavy atom, and L3 is the COM of an inhibitor heavy atom satisfying 30⁰ ≤ ∠L1L2L3 ≤150⁰ and the atoms bonded to the heavy atom. The atoms composing the six anchor points for Src-PP1 and Src-dasatinib were calculated according to the modeled X-ray structures and are as follows.

For c-Src-PP1: L1 (C5, N8, C9, C11 of PP1), L2 (C14, C15, C16, HC15 of PP1), L3 (C6, N10, H102, H101 of PP1), P1 (N, CA, C, O of Tyr82), P2 (N, CA, C, O of Glu81), P3 (N, CA, C, O of Val79).

For c-Src-dasatinib: L1 (C2, C, S of dasatinib), L2 (C3, N2, C4, HN2 of dasatinib), L3 (C14, H161, H153, H152 of dasatinib), P1 (N, CA, C, O of Tyr82), P2 (N, CA, C, O of Glu81), P3 (N, CA, C, O of Val79).

The Euler angles α (P1-L1-L2), β (P2-P1-L1-L2), and γ (P1-L1-L2-L3) define the orientation of the inhibitor with respect to the protein kinase and the polar angles θ (P2-P1-L1) and φ (P3-P2-P1-L1) define the position of the inhibitor with respect to the kinase.

**Obtaining structural ensembles from clustering**

*k*-means clustering analysis was performed using the analysis tools in the GENESIS^4, 5^ MD program on replicas at 310 K, separately for each of the bound, intermediate and encounter regions. Frames used for analysis were separated by 105 ps. Heavy atoms of the kinase and the inhibitors were used for fitting, heavy atoms of the inhibitor and of residues 1-150 of c-Src were used for clustering. The number of output clusters was n = 10. After the initial clustering, representative structures of the ten clusters were placed on the two-dimensional (2D) free-energy landscape and clusters residing in the same energetic minima were assigned into a single group, which defined the groups B1, B2, I1-I4, E1-E7 for Src-PP1 (“forward”) and B1, B2, I1-I3, and E1-E5 for Src-dasatinib (“forward”) in Figure 4 in the main text.

**Temperature replica exchange MD (T-REMD)^6^ simulations of dasatinib in water**

For the T-REMD simulations of dasatinib in solution, only the dasatinib inhibitor was extracted from the bound form (used for the gREST/REUS simulations) and solvated with 5,451 water molecules in a box of dimensions 53 × 60 × 54. The system was minimized for 1,000 steps while restraining the heavy atoms of dasatinib. Then, it was heated to 310 K during 100 ps under NVT conditions, followed by a 100 ps equilibration under NPT conditions using the Langevin thermostat and barostat. Finally, the system was equilibrated for 6 ns under NPT conditions with all restraints removed.

For the T-REMD simulations, 64 replicas covering the temperature range of 310-497 K were prepared and equilibrated under NVT conditions for 1 ns without any exchange attempts. The production run for T-REMD was performed for a total of 400 ns per replica with exchange attempts every 2 ps. The leapfrog integrator with a timestep of 2 fs was used throughout the equilibrations and production stages of the T-REMD simulations. All other parameters were identical to those used for the gREST/REUS simulations.

| **Table S1.** gREST/REUS simulation parameters | | | | |
| --- | --- | --- | --- | --- |
| System | Solute temperatures, K | gREST solute region protein residues | REUS replica distance, Å | REUS replicas force constants, kcal/mol/Å^2^ |
| Src-PP1 (“forward”) | 310, 344, 382, 426, 478, 539, 609, 692  310, 344, 382, 426, 478, 539, 609, 692 | 15, 23, 35, 37, 78, 80, 81, 83, 85, 87, 135, 146 | 3.00, 3.60, 4.00, 4.30, 4.60, 4.90, 5.20, 5.50, 5.80, 6.10, 6.60, 7.10, 7.70, 8.30, 8.90, 9.50, 10.10, 10.70, 11.30, 11.90, 12.50, 13.10, 13.70, 14.30, 14.90, 15.50, 16.10, 16.70, 17.30, 17.90 | 2.0, 2.0, 4.0, 4.0, 4.0, 4.0, 4.0, 3.0, 2.0, 2.0, 2.0, 2.0, 2.0, 2.0, 2.0, 2.0, 2.0, 2.0, 2.0, 2.0, 2.0, 2.0, 2.0, 2.0, 2.0, 2.0, 2.0, 2.0, 2.0, 2.0 |
| Src-PP1 (“reverse”) |  |  | 3.00, 3.20, 3.40, 3.60, 3.80, 4.10, 4.60, 5.10, 5.60, 6.15, 6.70, 7.25, 7.80, 8.20, 8.60, 9.10, 9.65, 10.20, 10.75, 11.35, 12.00, 12.65, 13.30, 13.95, 14.60, 15.25, 15.95, 16.65, 17.35, 18.05 | 4.0, 4.0, 4.0, 4.0, 4.0, 4.0, 3.0, 3.0, 3.0, 3.0, 3.0, 3.0, 3.0, 2.0, 2.0, 2.0, 2.0, 2.0, 2.0, 2.0, 2.0, 2.0, 2.0, 2.0, 2.0, 2.0, 2.0, 2.0, 2.0, 2.0 |
| Src-dasatinib (“forward”) | 310, 343, 381, 423, 471, 528, 590, 663  310, 343, 381, 423, 471, 528, 590, 663 |  | 3.00, 3.90, 4.80, 5.30, 5.80, 6.30, 6.90, 7.50, 8.20, 9.00, 9.80, 10.50, 11.20, 11.90, 12.60, 13.30, 14.00, 14.70, 15.40, 16.10, 16.80, 17.50, 18.20, 18.90, 19.60, 20.30, 21.00, 21.70, 22.40, 23.10 | 2.0, 2.0, 2.0, 2.0, 2.0, 2.0, 2.0, 2.0, 2.0, 2.0, 2.0, 2.0, 2.0, 2.0, 2.0, 2.0, 2.0, 2.0, 2.0, 2.0, 2.0, 2.0, 2.0, 2.0, 2.0, 2.0, 2.0, 2.0, 2.0, 2.0 |
| Src-dasatinib (“reverse”) |  |  | 3.0, 3.60, 4.20, 4.80, 5.40, 6.00, 6.60, 7.20, 7.90, 8.60, 9.30, 10.00, 10.70, 11.40, 12.10, 12.80, 13.50, 14.20, 14.90, 15.60, 16.30, 17.00, 17.70, 18.40, 19.15, 19.90, 20.65, 21.40, 22.20, 23.00 | 4.0, 4.0, 4.0, 4.0, 4.0, 4.0, 4.0, 4.0, 3.0, 2.0, 2.0, 2.0, 2.0, 2.0, 2.0, 2.0, 2.0, 2.0, 2.0, 2.0, 2.0, 2.0, 2.0, 2.0, 2.0, 2.0, 2.0, 2.0, 2.0, 2.0 |

| **Table S2.** Kinase-inhibitor nonbonded interaction probabilities (residue number^1^/contact probability^2^) | | | | | | | | | | | |
| --- | --- | --- | --- | --- | --- | --- | --- | --- | --- | --- | --- |
| Src-PP1 (“forward”) | | | | | | Src-dasatinib (“forward”) | | | | | |
| Bound | | Intermediate | | Encounter | | Bound | | Intermediate | | Encounter | |
| 83 | 86 | 15 | 66 | 33 | 19 | 80 | 87 | 15 | 72 | 20 | 32 |
| 35 | 74 | 83 | 61 | 82 | 18 | 83 | 86 | 84 | 56 | 21 | 29 |
| 81 | 72 | 86 | 57 | 17 | 17 | 35 | 81 | 82 | 52 | 18 | 27 |
| 80 | 67 | 84 | 47 | 84 | 16 | 37 | 80 | 86 | 45 | 19 | 24 |
| 135 | 63 | 82 | 43 | 21 | 15 | 86 | 77 | 17 | 39 | 22 | 24 |
| 15 | 55 | 23 | 34 | 37 | 14 | 82 | 76 | 83 | 38 | 17 | 18 |
| 37 | 49 | 90 | 25 | 20 | 13 | 84 | 72 | 33 | 37 | 37 | 17 |
| 82 | 47 | 85 | 22 | 23 | 13 | 15 | 69 | 90 | 33 | 38 | 16 |
| 23 | 37 | 135 | 21 | 25 | 12 | 135 | 66 | 25 | 32 | 49 | 15 |
| 86 | 30 | 87 | 20 | 15 | 12 | 36 | 59 | 23 | 31 | 44 | 15 |
| 87 | 29 | 35 | 14 | 68 | 11 | 52 | 48 | 87 | 30 | 48 | 12 |
| 52 | 21 | 25 | 11 | 85 | 11 | 23 | 44 | 85 | 28 | 51 | 11 |
| 78 | 15 | 17 | 10 | 18 | 10 | 145 | 39 | 16 | 28 | 68 | 10 |
|  |  |  |  |  |  | 78 | 32 | 14 | 24 |  |  |
|  |  |  |  |  |  | 56 | 26 | 37 | 16 |  |  |
|  |  |  |  |  |  | 85 | 25 | 21 | 16 |  |  |
|  |  |  |  |  |  | 81 | 22 | 18 | 14 |  |  |
|  |  |  |  |  |  | 146 | 20 | 20 | 14 |  |  |
|  |  |  |  |  |  | 65 | 13 | 13 | 12 |  |  |
|  |  |  |  |  |  |  |  | 132 | 12 |  |  |
|  |  |  |  |  |  |  |  | 22 | 12 |  |  |
| ^1^Left column for each region displays residue number.  ^2^Right column for each region displays contact probability (in %) for the residue. Residues with contact probabilities of >10% are displayed. | | | | | | | | | | | |

| **Table S3.** Kinase-inhibitor hydrogen bond probabilities (residue number^1^/contact probability^2^) | | | | | | | | | | | |
| --- | --- | --- | --- | --- | --- | --- | --- | --- | --- | --- | --- |
| Src-PP1 (“forward”) | | | | | | Src-dasatinib (“forward”) | | | | | |
| Bound | | Intermediate | | Encounter | | Bound | | Intermediate | | Encounter | |
| 83 | 77 | 83 | 24 | 33 | 6 | 83 | 82 | 33 | 26 | 20 | 19 |
| 81 | 59 | 82 | 6 | 37 | 5 | 80 | 46 | 15 | 21 | 21 | 5 |
| 80 | 8 | 90 | 6 |  |  |  |  | 90 | 19 |  |  |
|  |  | 15 | 5 |  |  |  |  | 84 | 15 |  |  |
|  |  |  |  |  |  |  |  | 83 | 14 |  |  |
|  |  |  |  |  |  |  |  | 87 | 9 |  |  |
|  |  |  |  |  |  |  |  | 82 | 7 |  |  |
|  |  |  |  |  |  |  |  | 146 | 5 |  |  |
|  |  |  |  |  |  |  |  | 17 | 5 |  |  |
| ^1^Left column for each region displays residue number.  ^2^Right column for each region displays contact probability (in %) for the residue. Residues with contact probabilities of >5% are displayed. | | | | | | | | | | | |

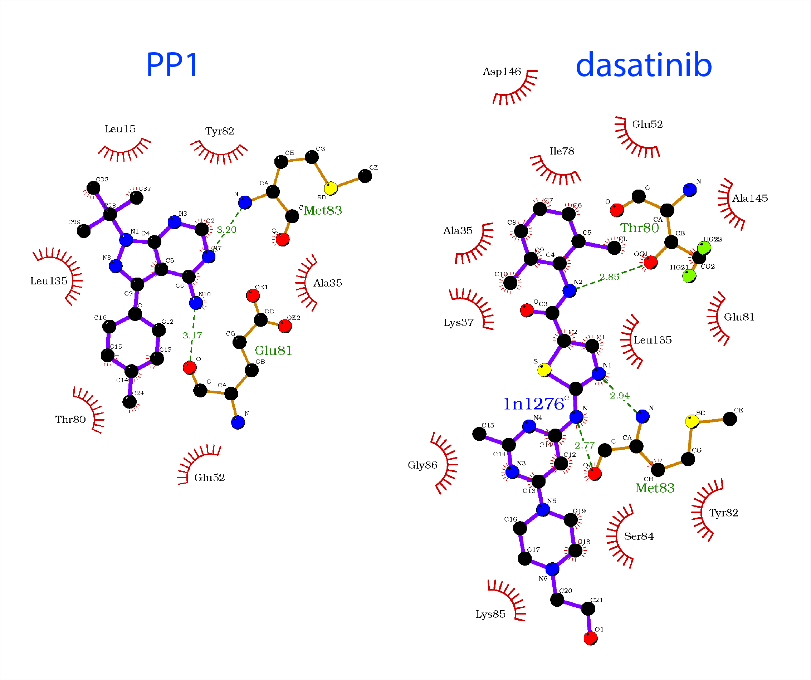

**Figure S1.** Nonbonded contacts and hydrogen bonds for PP1 (left) and dasatinib (right) according to the modeled X-ray structure (1Y57^7^/1QCF^8^ for c-Src-PP1, 1Y57^7^/3G5D^9^ for c-Src-dasatinib), created by ligplot+^10^.

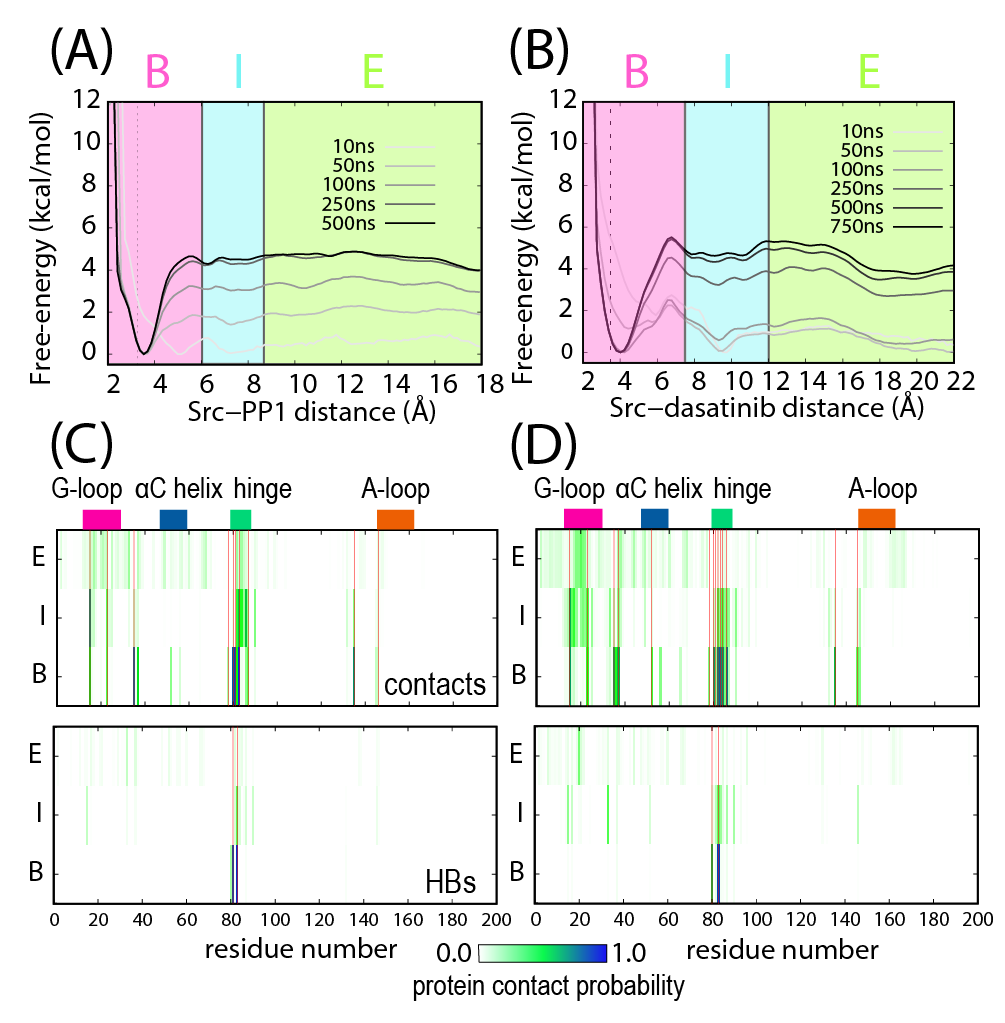

**Figure S2.** (A), (B) Free-energy profiles at 310 K along the c‑Src‑dasatinib (A) and c-Src-PP1 (B) distance for the “reverse” simulations. The kinase-inhibitor distances in the modeled X-ray structures (3.25 Å, PDB 1Y57^7^/1QCF^8^ for PP1 and 3.48 Å, PDB 1Y57^7^/3G5D^9^ for dasatinib) are shown in dotted line. Division between regions are marked in solid lines. (C), (D) Kinase-inhibitor contact (top) and hydrogen bond (HB, bottom) probabilities for c-Src residues 1-200 at 310K for “reverse” simulations for PP1 (C) and dasatinib (D). Location of binding site residues are shown in red vertical lines.

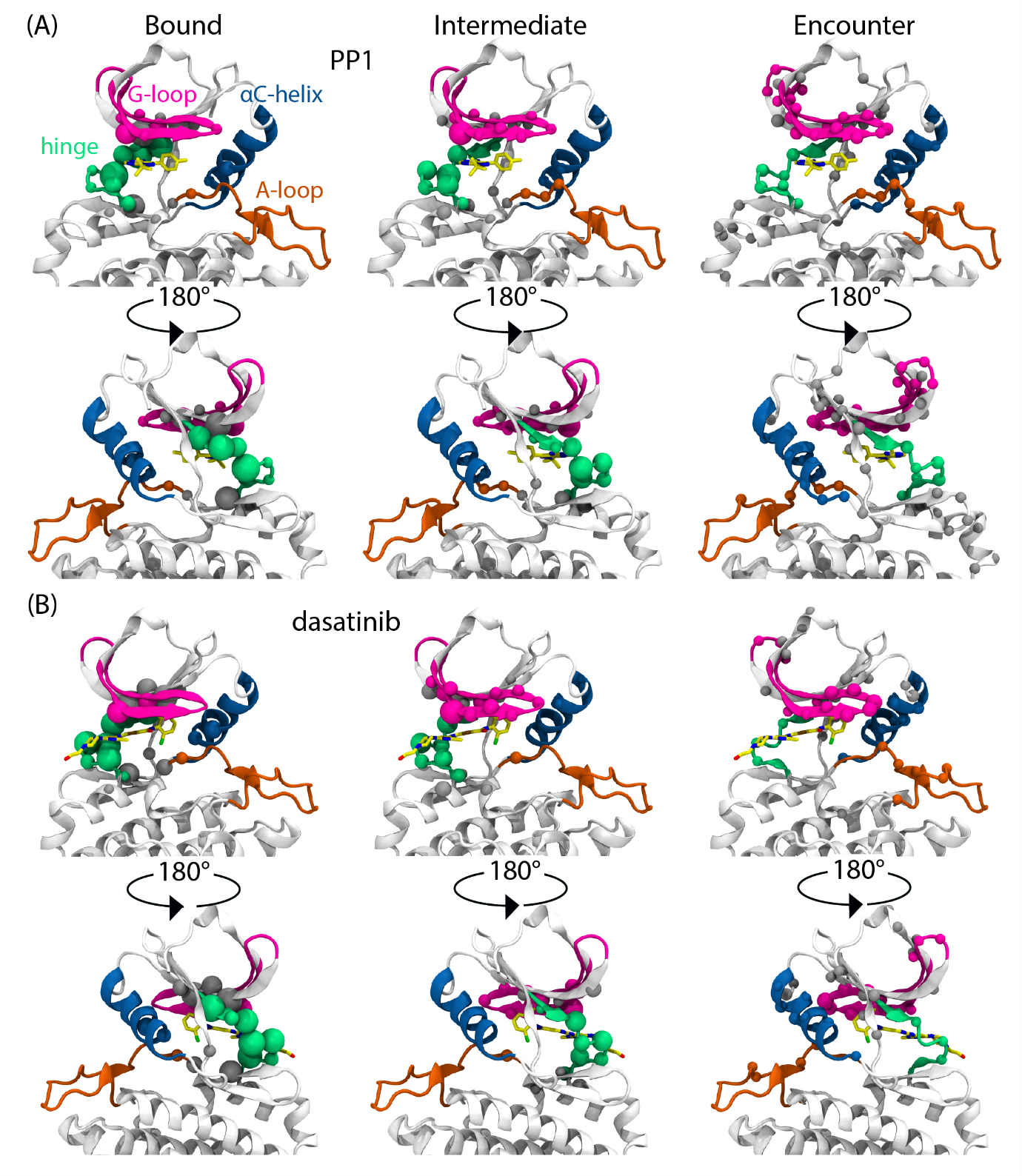

**Figure S3.** Interacting protein residues in the bound, intermediate and encounter regions for Src-PP1 (“forward”) (A) and Src-dasatinib (“forward”) (B), shown on the modeled X-ray structure of the respective complexes, from the front (top) and back (bottom) views. Residues with contact probabilities larger than 5% are shown as spheres of sizes proportional to the contact probability. Residues belonging to important structural motifs are colored.

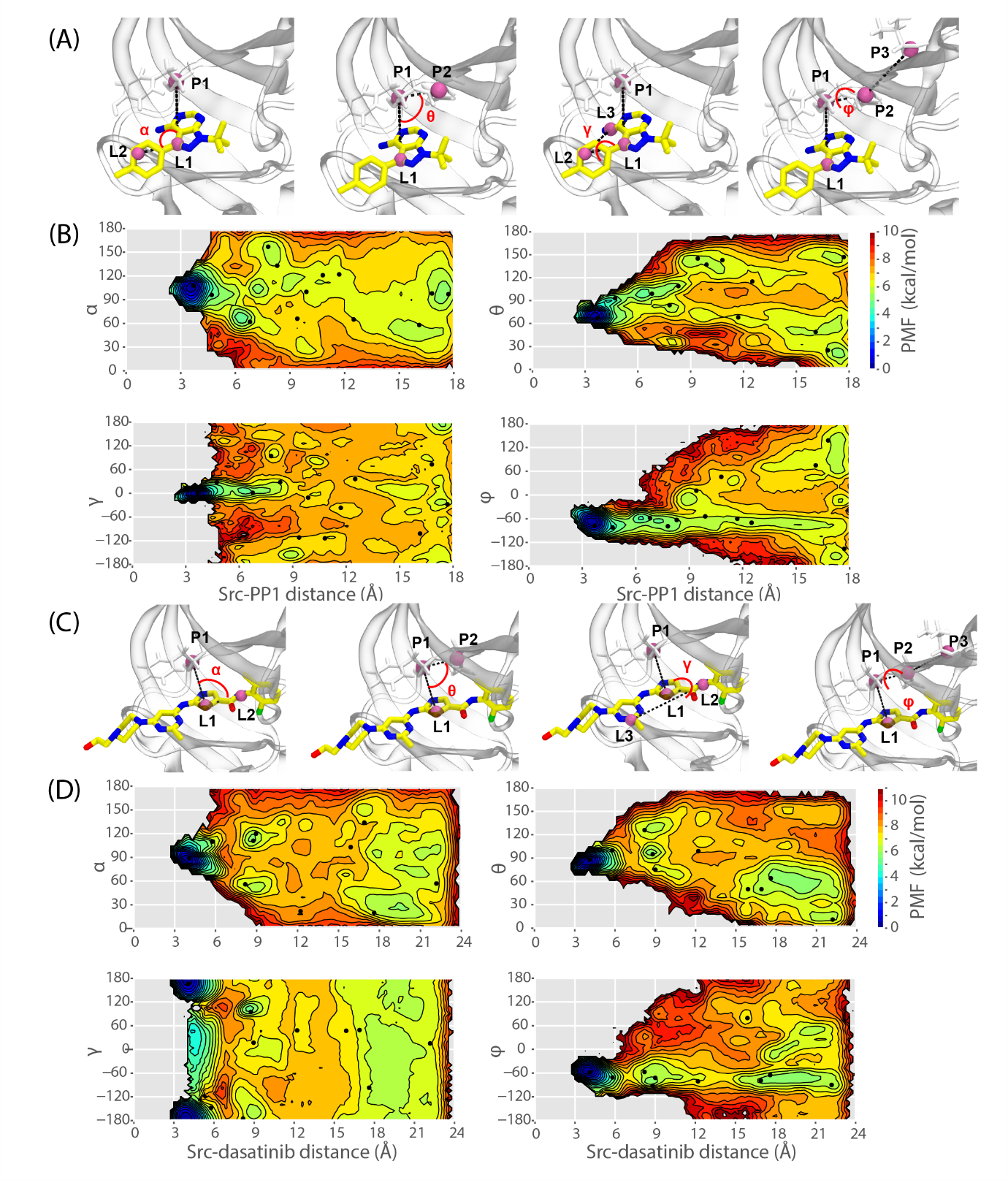

**Figure S4.** (A), (C) Definitions of Euler angles α and γ and polar angles θ and φ for defining the inhibitor orientation with respect to the kinase active site for c-Src-PP1 (A) and c-Src-dasatinib (C). (B), (D) The 2D free-energy landscapes at 310 K along the c-Src-PP1 (B) and c-Src-dasatinib (D) distances and the positional and orientational angles for the “forward” simulations at 310 K. Location of representative structures are marked as black dots on the plots.

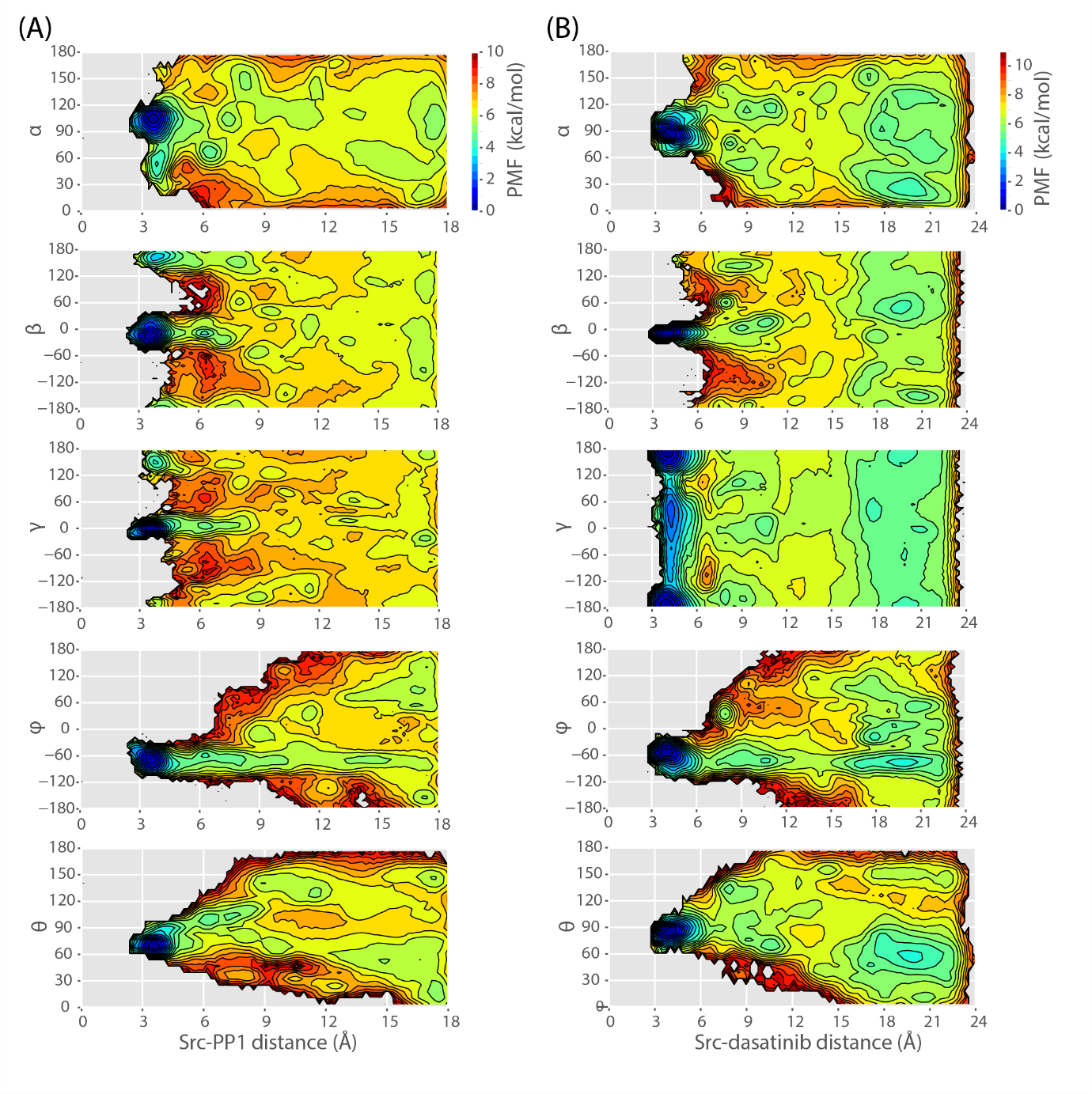

**Figure S5.** 2D free-energy landscapes at 310 K along the c-Src-PP1 (A) and c-Src-dasatinib (B) distances and the positional and orientational angles for the “reverse” simulations.

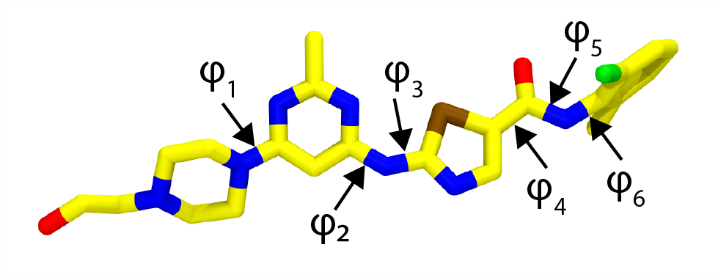

**Figure S6.** The six dihedral angles for dasatinib.

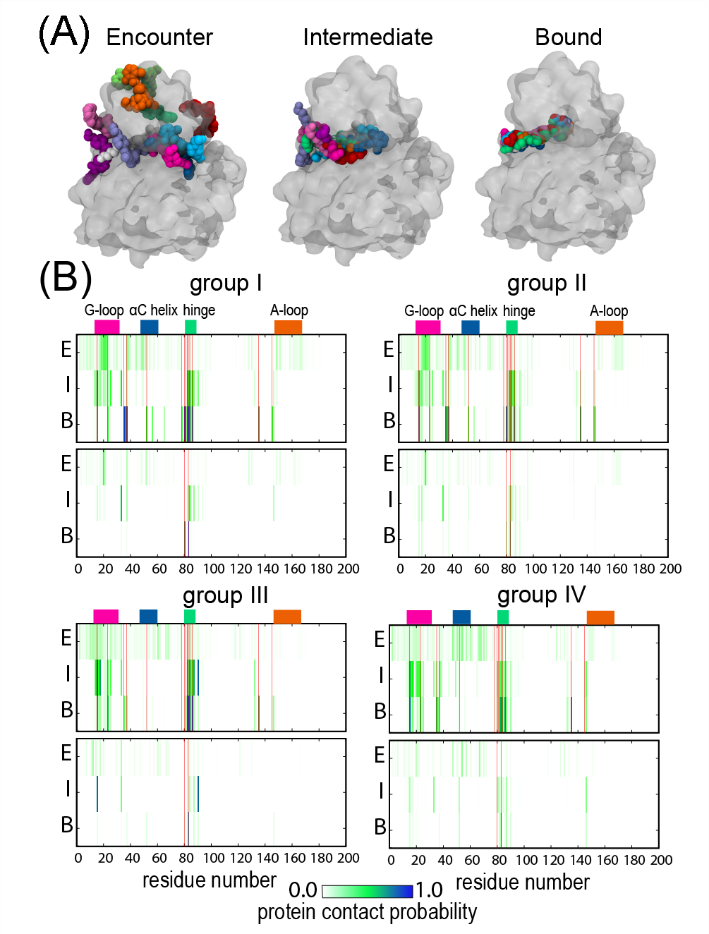

**Figure S7.** (A) Dominant inhibitor conformations with respect to the protein kinase for the encounter, intermediate and the bound regions, extracted from *k*-means clustering. (B) Kinase-inhibitor nonbonded contact (top of each panel) and HB (bottom) probabilities for c-Src residues 1-200 at 310K for Src-dasatinib (“forward”) simulations, divided into inhibitor conformers I-IV. Location of binding site residues are shown in red vertical lines.
